# Neutrophil subsets in SLE exhibit increased glycolysis that correlates with disease activity

**DOI:** 10.64898/2026.05.14.725124

**Authors:** Anjali S. Yennemadi, Natasha Jordan, Sophie Diong, Faye K Murphy, Sarah Quidwai, Mark A Little, Joseph Keane, Gina Leisching

## Abstract

Systemic lupus erythematosus (SLE) is a chronic autoimmune disease characterised by sustained type I interferon signalling and widespread immune dysregulation. Low-density neutrophils (LDNs) are expanded in SLE and display pro-inflammatory and tissue-damaging properties. However, their metabolic phenotype remains poorly defined. Here, we performed a comprehensive metabolic characterisation of circulating LDNs and normal-density neutrophils (NDNs) from patients with SLE and matched healthy individuals (HC).

Neutrophil subsets were isolated from peripheral blood of SLE patients and HC donors using a two-step protocol of negative selection and Percoll density centrifugation. Immunophenotyping phenotype was carried out by flow cytometry to assess phenotypic expression of common neutrophil markers CD15, CD16, CD10, CD66b, CD62L, MPO, and IL-1β. Bioenergetic profiling of LDNs and NDNs was performed *in situ* using the Seahorse MitoStress test to measure oxygen consumption rate (OCR) and extracellular acidification rate (ECAR). Metabolic flexibility and phenotypic alterations were assessed in LDNs and NDNs following inhibiting mitochondrial metabolism with oligomycin and glycolysis with 2DG.

We found that SLE LDNs exhibit an immature phenotype compared with autologous and healthy NDNs, as determined transcriptionally by *C/EBPε* and by surface protein expression levels of CD10. Both LDNs and NDNs from SLEDAI≥4 patients demonstrated significantly elevated ECAR relative to HC neutrophils. Further, SLE LDNs displayed enhanced metabolic flexibility, with the capacity to switch towards a glycolytic phenotype under metabolic stress conditions. Inhibition of glycolysis altered the inflammatory and maturation-associated phenotype of both SLE neutrophil subsets, indicating a direct link between cellular metabolism and pathogenic neutrophil function.

Collectively, these findings identify fundamental metabolic alterations in SLE neutrophil subsets and support neutrophil immunometabolism as a potential therapeutic target in SLE.

## Introduction

Systemic lupus erythematosus (SLE) is a chronic autoimmune disease that is distinguished by the sustained upregulation of interferon (IFN)α and people with SLE exhibit pathological manifestations that afflict multiple organ systems that are associated with disease activity. In recent years, aberrant immune cell behaviour in SLE has been dissected, however there is still an appreciable gap in our understanding of the role of neutrophil subsets and their contribution to the pathogenesis of this disease.

Neutrophil subsets, in particular low-density neutrophils (LDNs), are elevated in the circulation of SLE patients ^1–3^. LDNs from SLE patients have been identified as a circulating source of IFNα, as well as having damaging effects at the tissue level and on the vasculature.^1^ These circulating LDNs in SLE patients have been shown to display a pro-inflammatory phenotype and the ability to infiltrate sites of skin lesions. LDNs from skin biopsies of patients with SLE, discoid lupus, acute cutaneous lupus, subacute cutaneous lupus and lupus panniculitis have also been shown to have a higher capacity for producing neutrophil extracellular traps (NETs) compared to healthy donors at the sites of these lesions^1,4^. Additionally, LDNs possess distinct cellular biomechanics which enhance retention in the microvasculature, contributing to endothelial damage^5^. The pro-inflammatory nature of SLE LDNs has also been characterised by augmented secretion of other pro-inflammatory cytokines such as TNF and IFNγ^1^, with TNF being associated with worsening kidney histological activity in lupus nephritis patients^6^.

Previously, we have proposed that the pathogenic alterations at sites of tissue damage can be attributed, at least in part, to the dysfunctional metabolic activities of immune cells in this niche, particularly LDNs, as a result of the chronic exposure to type I IFNs^7^. Thus, their involvement in causing inflammation and damage at tissue sites and the mechanisms driving this pathogenic phenotype should be investigated further.

A recent study of otherwise healthy granulocyte colony-stimulating factor (G-CSF)-treated donors found that LDNs had enhanced mitochondrial bioenergetic capacity compared to normal density neutrophils (NDN) ^3^. Although the authors suggest that the observed metabolic features are similar to what would be expected in SLE patients, no direct bioenergetic comparison of SLE LDN or NDNs were conducted. The metabolic analyses also did not include the use of non-GCSF-treated (“healthy”) LDNs and NDNs as subset-specific comparators. Another in-depth study on LDNs from SLE patients also did not analyse LDNs from healthy donors,^8^ attributing this to their rarity. However, LDNs are readily accessible in healthy individuals (HC), as we and others have shown before^9–11^. Further, moving away from the traditional view of neutrophil metabolism being exclusively dependent on glycolysis, neutrophils have been shown to switch between glycolysis, OXPHOS, glycogenolysis, and fatty acid oxidation based on their maturity, nutrient availability, and activation status^12^. This implies that neutrophils are metabolically flexible depending on their environment, and positions neutrophil immunometabolism as a potentially druggable target that could be leveraged to improve therapeutic outcomes. However, owing to the lack of metabolism-based studies on neutrophil subsets, particularly in SLE patients, the effect of metabolic inhibitors on the phenotype of SLE LDNs and NDNs is not very well known.

The current study addresses these shortcomings with the aim of characterising the metabolism of LDNs and NDNs from SLE patients and HCs and is the first study showing fundamental metabolic differences that are directly attributable to SLE disease. We show that i) SLE LDNs are immature compared to both, healthy and autologous NDNs, ii) LDNs and NDNs from patients with more severe SLE have significantly higher glycolytic activity than HC neutrophil subsets, iii) LDNs from patients with more severe SLE are also able to switch to a glycolytic phenotype under conditions of metabolic stress, and iv) glycolysis alters the phenotype of both LDNs and NDNs from SLE patients.

## Materials and Methods

### Participant Recruitment

We recruited SLE patients attending routine clinical assessments at the Rheumatology Clinic at St. James’s Hospital. Patients fulfilled the 2019 EULAR/ACR Classification Criteria for Systemic Lupus Erythematosus.^13^ Peripheral blood was collected from SLE patients and age- and sex-matched healthy controls (HC) (Table 1) under full ethical approval for the sampling and processing of human biological samples from the St. James’s Hospital Research ethics committee. All participants involved in this study provided informed, written consent.

**Table 1:** Patient Characteristics and Donor Demographics.

| Characteristics | SLEDAI <4<br>(n = 15) | SLEDAI ≥4<br>(n = 20) | SLE combined<br>(n = 35) | Healthy<br>(n = 26) |
| --- | --- | --- | --- | --- |
| **Demographics** |  |  |  |  |
| Age, years (median, range) | 40 (17 - 62) | 32 (16 - 62) | 38 (16 - 62) | 33.5 (22 - 62) |
| Sex, n (%) |  |  |  |  |
| Female | 12 (80) | 18 (90) | 30 (85.7) | 17 (65.4) |
| Male | 3 (20) | 2 (10) | 5 (14.3) | 9 (34.6) |
| **Lupus History** |  |  |  |  |
| SLEDAI (median, range) | 0 (0 – 2) | 8 (4 – 22) | 4 (0 – 22) | - |
| ESR, mm/hour (median, range) | 10 (2 - 74) | 15 (2 - 49) | 13.5 (2 - 74) | - |
| ANC, x 10<sup>9</sup> cells/L (median, range) | 3.2 (1.8 - 6.2) | 3.1 (1.1 - 7.4) | 3.1 (1.1 - 7.4) | - |
| Low C3, n (%) | 0 | 5 (33.4) | 5 (14.3) | - |
| Low C4, n (%) | 1 (6.7) | 12 (80) | 13 (37.1) | - |
| **Autoantibodies, n (%)** |  |  |  |  |
| Anti-Ro | 8 (53.4) | 14 (70) | 22 (62.9) | - |
| Anti-Sm | 1 (6.7) | 2 (10) | 3 (8.6) | - |
| Anti-dsDNA | 2 (13.4) | 5 (33.4) | 14 (40) | - |
| ANA | 12 (80) | 20 (100) | 32 (91.4) | - |
| RNP | 1 (6.7) | 7 (35) | 8 (22.9) | - |
| Anti-phospholipid | 3 (20) | 6 (30) | 9 (25.7) | - |
| **Medication, n (%)** |  |  |  |  |
| Prednisolone | 6 (40) | 10 (50) | 16 (45.7) | - |
| <10mg daily | 3 (20) | 3 (15) | 6 (17.1) | - |
| ≥10 mg daily | 2 (13.4) | 3 (15) | 5 (14.3) | - |
| NAV | 1 (6.7) | 4 (20) | 5 (14.3) | - |
| Hydroxychloroquine | 13 (86.7) | 15 (75) | 28 (80) | - |
| Mycophenolate mofetil | 1 (6.7) | 7 (35) | 8 (22.9) | - |
| Azathioprine | 4 (26.7) | 3 (15) | 7 (18.9) | - |
| Mepacrine | 0 | 5 (33.4) | 5 (13.5) | - |
| Methotrexate | 2 (13.4) | 2 (10) | 4 (11.4) | - |
| Betamethasone valerate | 0 | 1 (5) | 1 (2.7) | - |
| Tacrolimus | 0 | 1 (5) | 1 (2.7) | - |
| Rituximab | 1 (6.7) | 0 | 1 (2.7) | - |
| Warfarin | 0 | 1 (5) | 1 (2.7) | - |
| Aspirin | 0 | 1 (5) | 1 (2.7) | - |
| Valsartan | 0 | 1 (5) | 1 (2.7) | - |
| Nil | 1 (6.7) | 0 | 1 (2.7) | - |

*SLEDAI: SLE Disease Activity Index; ESR: Erythrocyte sedimentation rate; ANC: Absolute neutrophil count; ANA: Antinuclear antibody; RNP: Ribonucleoprotein; NAV: Not available*

### LDN and NDN Isolation

Peripheral venous blood was collected from HC and SLE patients in Lithium-heparin vacuum containers. Total neutrophils were isolated from 12 mL blood within 2 h of phlebotomy using the EasySep™ Direct Human Neutrophil Isolation Kit as per manufacturer’s instructions (Stemcell Technologies). A negative selection protocol was employed to avoid activating the cells. LDNs and NDNs were then sorted using a discontinuous Percoll density gradient approach as we have previously described.^11^

### Flow Cytometry, Glycolysis Inhibition, and SPICE Analysis

For flow cytometry analysis, 1 × 10^5^ cells per condition were washed with PBS and centrifuged at 400 × g for 5 min at room temperature (RT) immediately after isolation. LDNs and NDNs were then resuspended in PBS and incubated with 2-deoxy-D-glucose (100mM; 2DG) to inhibit glycolysis or with PBS as an untreated control (‘Co’) for 15 min at 37°C. After 15 min, cells were washed with PBS and centrifuged as before.

LDNs and NDNs were then surface stained for flow cytometry using fluorochrome-conjugated antibodies against CD10, CD15, CD16, CD66b, CD62L (all 1:100), Fc block (1:50), and viability stain (1:200) in PBS for 20 min in the dark at RT. The cells were then washed and centrifuged as before, and fixed and permeabilized using the Foxp3 staining kit (eBioscience) as per manufacturer’s instructions. Cells were incubated for 30 min at 4°C with anti-MPO (1:200) and anti-IL-1β (1:100), for intra-cellular staining and then acquired on a Northern Lights spectral flow cytometer (Cytek Biosciences). See Table 2 for more details on the panel used. For all flow cytometry analyses, LDNs and NDNs were defined as CD15+ cells with high side scatter (CD15+ SSChi), and the frequency of sub-populations was reported as a percentage of total LDNs or NDNs, respectively.

**Table 2:** Antibodies, clones, and dilutions utilised for immunophenotyping by flow cytometry.

| Antigen | Clone | Fluorochrome | Manufacturer | Catalogue Code | Dilution |
| --- | --- | --- | --- | --- | --- |
| CD10 | HI10a | BV421 | Biolegend | 312218 | 1:100 |
| CD15 | W6D3 | PE/Dazzle™ 594 | Biolegend | 323038 | 1:100 |
| CD16 | 3G8 | FITC | Biolegend | 302005 | 1:100 |
| CD62L | DREG-56 | BV650 | Biolegend | 304831 | 1:100 |
| CD66b | G10F5 | PE/Cy7 | Biolegend | 305115 | 1:100 |
| IL1β | JK1B-1 | AlexaFluor647 | Biolegend | 508207 | 1:100 |
| MPO | 2C7 | PE | BioRad | MCA1757PE | 1:100 |
| FcR Blocking Reagent | N/A | N/A | Miltenyi Biotec | 130-059-901 | 1:50 |
| Zombie Aqua Fixable Viability Dye | N/A | N/A | Biolegend | 423102 | 1:200 |

Data were analysed using FlowJo Software (v10.10; BD Life Sciences). Simplified Presentation of Incredibly Complex Evaluations (SPICE) analysis was carried out on sub-population frequency data using Pestle (v2) and SPICE (v6.1) software after pre-processing data using FlowJo as per published guidelines ^14^.

### In situ Metabolic Analysis

LDNs and NDNs were seeded at 1.25 × 10^5^ cells/well in triplicate into a Seahorse XFp Cell Culture Microplate (Agilent Technologies) immediately after isolation and centrifuged for 1 min at 1000 RPM, followed by deceleration without braking. Seahorse XF RPMI was supplemented with 4.5 g/L d-glucose, 2 mM glutamine, 100 mM pyruvate and subsequently added to the cells. Following incubation in a CO_2_-free incubator at 37 °C for 60 min, the Agilent Cell MitoStress assay was performed. For this, basal oxygen consumption rate (OCR) and extra-cellular acidification rate (ECAR) were recorded, followed by sequential addition of 2 µg/ml oligomycin (Oligo), 0.5 µM carbonyl cyanide 4-(trifluoromethoxy) phenylhydrazone (FCCP) and 1 µM rotenone/antimycin A (Rot/AA). OCR and ECAR values were normalised using the Crystal Violet dye extraction growth assay and the Wave Desktop v2.6 Software (https://www.agilent.com) was used for analysis.

### In situ ATPase Inhibition in LDNs

LDNs were seeded into a flat-bottom 96 well plate at 1.25 × 10^5^ cells/well and centrifuged at 400 x g for 5 min immediately after isolation. Supernatants were discarded and LDNs were treated with either 2 µg/ml oligomycin (‘Oligo’) to inhibit mitochondrial ATPase (Complex V) activity, or with Seahorse XF RPMI supplemented as detailed above (‘untreated’) for 18 min to recapitulate the addition of oligomycin during the Agilent Cell MitoStress assay. The incubation was carried out in a CO_2_ incubator at 37°C, followed by addition of Seahorse XF RPMI to stop treatment, and centrifugation at 400 x g for 5 min. Supernatants were discarded and cells were lysed for RNA extraction as detailed below.

### qPCR

Total RNA was extracted using the RNeasy® Plus Mini Kit (Qiagen) according to manufacturer’s instructions. For cDNA synthesis, 0.1 µg RNA was converted to cDNA using the Quantitect® Reverse Transcription Kit (Qiagen). To ensure the removal of genomic DNA, “gDNA wipe-out buffer” was added to RNA (included in the kit) prior to the RNA conversion step. qPCR amplification was performed using a 384-well QuantStudio™ 5 Real-Time PCR System (Applied Biosystems) to assess differentially expressed genes *(CEBPA, CEBPD, CEBPE, CEBPG; HK2, PFKM, PKM2, PFKFB3)* using QuantiTect® primer assays at a reaction volume of 20 μl with reference gene *18S*. The amplification procedure entailed 45 cycles of 95°C for 10 sec followed by 60°C for 10 sec and finally 72°C for 10 sec. Gene expression was evaluated at baseline for *CEBPA, CEBPD, CEBPE, CEBPG* and so expression levels were calculated as 2^-ΔCT^, in keeping with recommended alterations to the 2^−ΔΔCT^ method ^15^. Fold change in expression levels for glycolysis genes *(HK2, PFKM, PKM2, PFKFB3)* was calibrated using the 2^−ΔΔCT^ method. All biological replicates were run in triplicate.

### Statistical analysis

Statistical analyses were performed using GraphPad Prism 11 software. All data were analysed as nonparametric. Statistically significant differences were determined using the Kruskal-Wallis test with post-hoc Dunn’s test or two-way ANOVA with post-hoc Fisher’s Least Significant Difference test, as appropriate. Where values were missing in a dataset for a two-way ANOVA, mixed-effects analysis with post-hoc Fisher’s Least Significant Difference test was performed. All correlation analyses were performed using Spearman’s rank correlation, and Spearman’s r values were reported, p-values of < 0.05 were considered statistically significant.

## Results

### LDNs are elevated, immature, display heterogeneity, and are phenotypically distinct in SLE

LDNs and NDNs were isolated from whole blood of SLE patients (n = 35) and healthy individuals (n = 26) using a two-step isolation process we have described previously^9^ (Table 1, Fig.1A). Following isolation, we noted that the number of total neutrophils recovered through our isolation protocol was lower than that from full blood counts. However, no difference was noted in the total neutrophil counts after isolation or from clinically obtained absolute neutrophil count (ANC) data between SLEDAI ≥4 and SLEDAI <4 patients (Fig.S1A). Further, we observed that patients with severe disease (SLEDAI ≥4) had significantly higher levels of LDNs per ml blood compared to HC (Fig.1B). Conversely, SLE patients with active disease also had significantly lower NDN counts compared to both, patients with inactive disease and HCs (Fig.1B). While corticosteroid use has been associated with neutrophilia, we found no significant correlation between clinically reported ANC and steroid dose (r= 0.4516, p= 0.1638; Fig.S1B).

**Figure 1:**
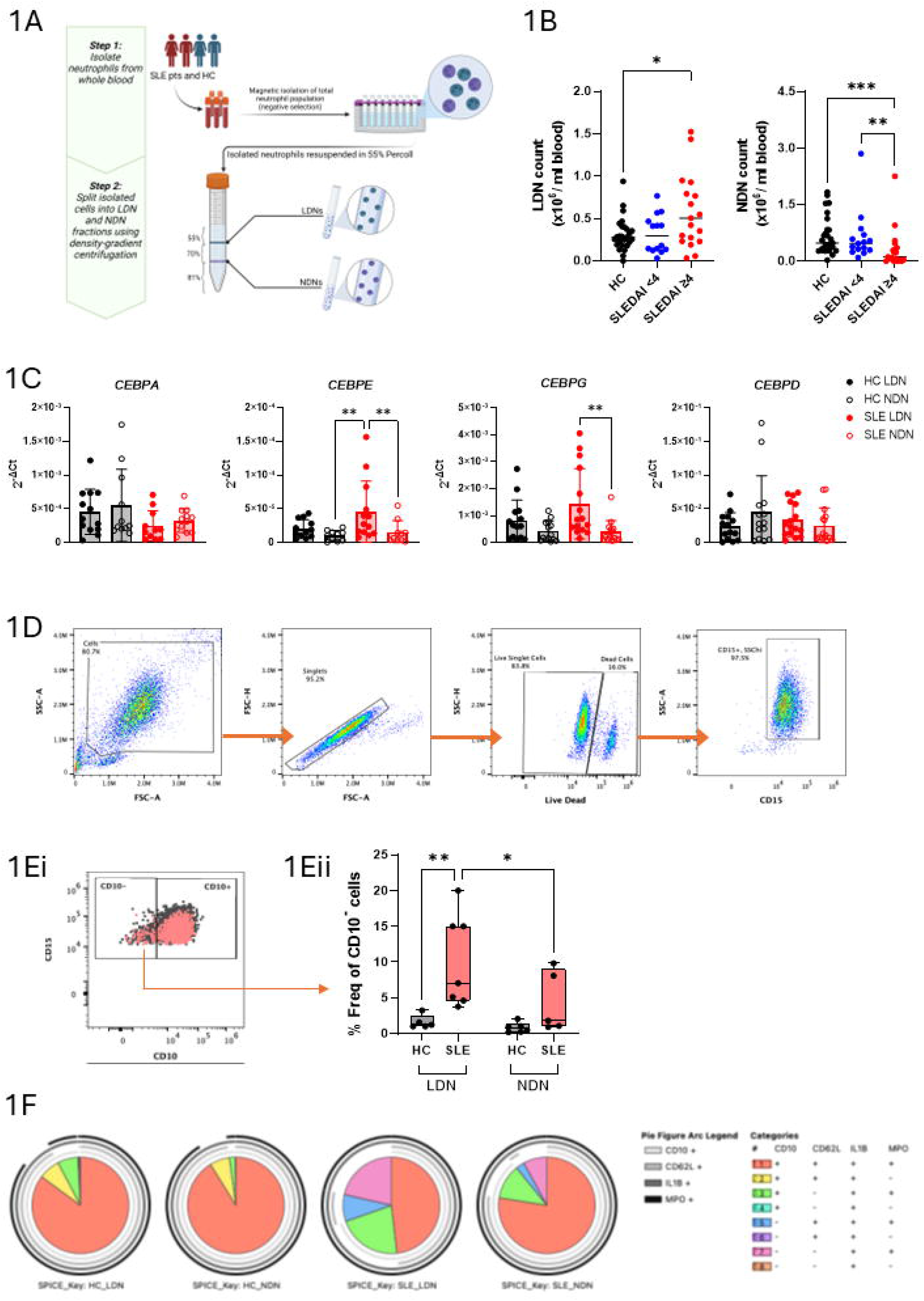
LDNs are expanded in circulation, immature, and heterogenous in SLE. (A) LDNs and NDNs were isolated from SLE and HC peripheral blood using a 2-step protocol. (B) SLE patients with high disease activity (SLEDAI ≥4) have elevated circulating LDNs and reduced NDNs per ml blood compared to healthy controls (HC). (C) qPCR assessment of neutrophil differentiation markers *C/EBP α, ε, γ, and δ* revealed that SLE LDNs express *C/EBPε* significantly higher than autologous NDNs and HC NDNs. (D) Flow cytometry gating strategy for isolated LDNs and NDNs: Debris, doublets, and dead cells were excluded. CD15+, SSChi cells were gated on and defined as neutrophils. (E) (i) Next, CD10 expression was used to discriminate between mature (CD10+) and immature (CD10-) populations, (ii) SLE patients have an increased frequency of immature (CD10-) LDNs compared to HC LDNs and autologous NDNs. (F) Pie charts generated following SPICE analysis subpopulation frequencies indicate heterogeneity in SLE LDNs driven by differences in CD10, CD62L, and MPO co-expression. *All analyses were Kruskal-Wallis tests, *p<0*,*05. SLEDAI: SLE Disease Activity Index; CEBP: CCAAT-enhancer binding protein; SPICE: Simplified Presentation of Incredibly Complex Evaluations*.

The CCAAT-enhancer binding protein (C/EBP) transcription factor family (α, β, γ, δ, ε, and ζ) has been described as essential for neutrophil differentiation. Of these proteins, C/EBPε is specifically required at the myelocyte and meta-myelocyte stages ^16–18^, and as such has been used to describe neutrophil maturity at a transcriptional level^19^. Thus, we quantified expression of transcription factors *C/EBPα, γ, ε*, and δ to assess maturity in LDNs and NDNs. We observed that LDNs from SLE patients express increased levels of *C/EBPε* compared to their autologous NDNs and compared to HC NDNs. However, the expression of *C/EBPα, γ*, and δ, remained comparable across cell types within and across groups (Fig.1C). These data suggest that SLE patients with active disease comprise a higher number of circulating LDNs compared to HC donors or inactive disease patients, and that these LDNs likely possess an immature phenotype corresponding to the myelocyte and meta-myelocyte stages, versus autologous NDNs.

We then ran flow cytometry to immunophenotype LDN and NDN sub-populations. SLE patients were assessed combined and compared to HC, as no significant differences were noted between LDNs and NDNs from SLEDAI ≥4 and SLEDAI <4 patients. First assessing potential differences in cell size and granularity, we found no statistically significant differences in FSC or SSC between LDNs and NDNs from both SLE patients and healthy individuals (Fig.S1C). Although there exists considerable debate in the literature with regards to whether LDNs are simply degranulated and therefore less-buoyant NDNs^20,21^, our data indicate that LDNs and NDNs harbour similar granularity and thus, warrant investigation as separate neutrophil subtypes.

A number of studies have utilised the differential expression of CD10 as a marker for maturity on neutrophils ^2,8,22^. In keeping with previous findings, we also observed two LDN sub-populations (Fig.1E(i)), CD10+ (mature) and CD10- (immature). Based on this dichotomy in CD10 expression, we found that SLE LDNs have a higher frequency of immature CD10-cells (Fig.1E(ii)) and lower frequency of mature CD10+ cells (Fig.S1D) compared with both, autologous NDNs and HC LDNs. On the other hand, the SLE NDN fraction showed comparable frequencies of both CD10 subsets compared to healthy donors (Fig. 1E(ii), S1D).

Immunophenotyping of SLE and HC neutrophil subsets also revealed phenotypic heterogeneity within each group. Pie charts generated using data from SPICE analysis (Table S1) show differences in subpopulation frequencies across HC LDNs (n=5), HC NDNs (n=5), SLE LDNs (n=5), and SLE NDNs (n=4). As LDNs and NDNs across both categories were CD15+, CD16+, and CD66b+, the analysis was focussed on the difference in frequencies of CD10, CD62L, IL-1β, and MPO co-expressing sub-populations. Strikingly, SLE LDNs appear to be the most heterogeneous group with the largest fractions of immature, activated cells (CD10-CD62L-IL-1β+MPO+; pink), followed by SLE NDNs which have a similar profile but with a comparatively lower proportion of immature, activated cells (pink; Fig.1F). In comparison with SLE, both HC LDNs and NDNs show a similar profile to each other of a small fraction of activate, mature cells (CD10+CD62L-IL-1B+MPO+; light green).

Together these data suggest that while LDNs are elevated only in individuals with active SLE, the SLE LDN population as a whole display an immature phenotype as well as an increased proportion of immature, activated cells when compared with SLE NDNs as well as LDNs from healthy individuals.

### NDNs from SLE patients with active disease display increased baseline glycolytic activity which correlates with disease severity

We performed the Agilent cell Mitostress assay to test mitochondrial fitness and function of LDNs and NDNs from SLE patients and healthy donors. Prior studies have shown that neutrophils do not ordinarily respond to metabolic inhibitors^23^. We were surprised to note that only NDNs from SLE patients with active disease responded to FCCP (a mitochondrial uncoupler) indicating that they can engage mitochondrial metabolism, in addition to glycolysis, in response to increased bioenergetic demands (Fig 2A). Further, NDNs from SLE patients, irrespective of disease activity, exhibit overall increased ECAR kinetics, a proxy measure for glycolysis (Fig 2B). Looking at the specific metabolic parameters, basal OCR of NDNs from SLE patients as a whole was not significantly different compared to healthy NDNs (Fig 2C). Similarly, OCR-dependent parameters of NDNs such as maximal respiration, spare respiratory capacity, non-mitochondrial oxygen consumption and ATP production were also at comparable levels across groups (Fig 2C). More specifically, SLE NDNs showed a significant increase in baseline glycolysis compared to NDNs from HC (Fig 2C). Additionally, when plotted against the SLEDAI score of the patients, this increase in ECAR showed a moderate positive correlation (r=0.5381, p = 0.012). Overall, the increase in baseline glycolysis of NDNs from patients with active disease drives them towards a relatively more energetic state compared to NDNs from patients with inactive disease or HC, as represented in the energy map (Fig 2E).

**Figure 2:**
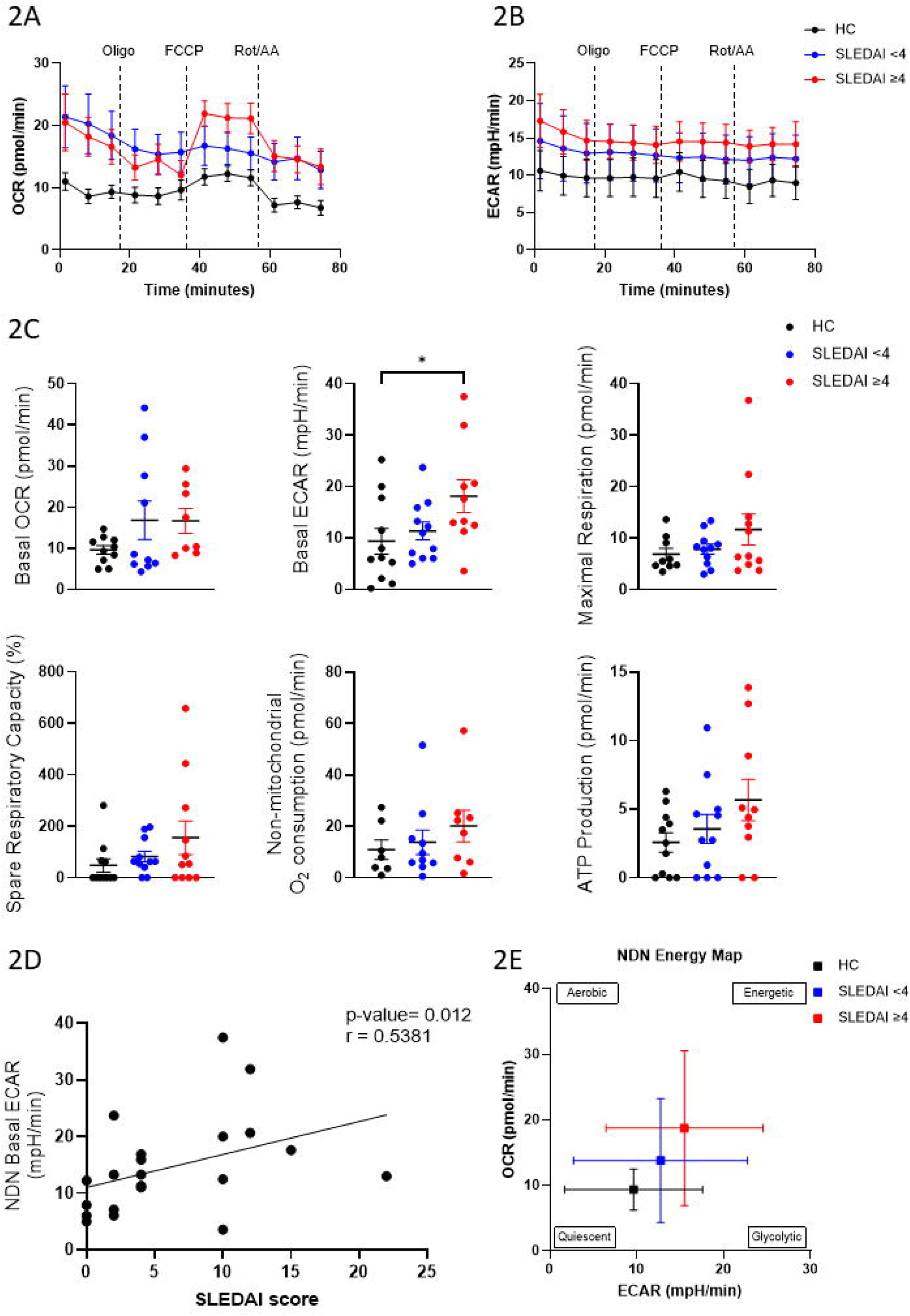
SLE NDNs are more glycolytic than HC NDNs, and this correlates with disease activity. (A) The Seahorse MitoStress test reveals that basal OCR kinetics are comparable across NDNs from SLE patients and HC. Of note, SLEDAI ≥4 NDNs exhibit sensitivity to FCCP indicating the use of mitochondrial respiration when bioenergetic demand is increased in this cohort. (B) NDNs from patients with SLEDAI ≥4 show higher basal ECAR kinetics compared to both SLEDAI<4 and HC NDNs. (C) This increased glycolysis is reflected in the basal ECAR graph as well. All other metabolic parameters have comparable values. (D) Plotting basal ECAR values against SLEDAI scores revealed a statistically significant, moderate positive correlation. (E) The energy map shows the more energetic profile of SLEDAI≥4 NDNs, relative to SLEDAI <4 NDNs and HC NDNs. *All graphs depict mean values with SEM, and all analyses were Kruskal-Wallis tests, *p<0*,*05. SLEDAI: SLE Disease Activity Index; OCR: Oxygen Consumption Rate; ECAR: Extracellular Acidification Rate; SEM: Standard Error of Mean*.

### LDNs from SLE patients display increased baseline glycolytic activity which correlates with disease severity

Similar to their autologous NDNs, LDNs from SLE patients also showed no differences in their overall OCR kinetics, irrespective of disease activity (Fig 3A). Interestingly however, LDNs from all patients and HCs responded to FCCP, indicating a prominent role for mitochondria in this neutrophil subtype. Despite this, LDNs from SLE patients with active disease exhibited a trend towards a decreased spare respiratory capacity compared to LDNs from HC and patients with inactive disease. LDNs from SLE patients as a whole displayed comparable levels of maximal respiration, spare respiratory capacity, non-mitochondrial oxygen consumption, and ATP production compared to HC LDNs (Fig 3C). Basal ECAR of SLE LDNs from patients with active disease was elevated compared to those from patients with inactive disease and HC LDNs as well (Fig 3B, C). This observation is reflected in the moderately high positive correlation between basal ECAR of LDNs from SLE patients with their SLEDAI scores (r = 0.5998, p = 0.002; Fig 3D). Overall, the elevated basal glycolysis observed in LDNs from patients with active disease shifts them into a more energetic state when compared with LDNs from patients with inactive disease or HC, as illustrated in the energy map (Fig 3E).

**Figure 3:**
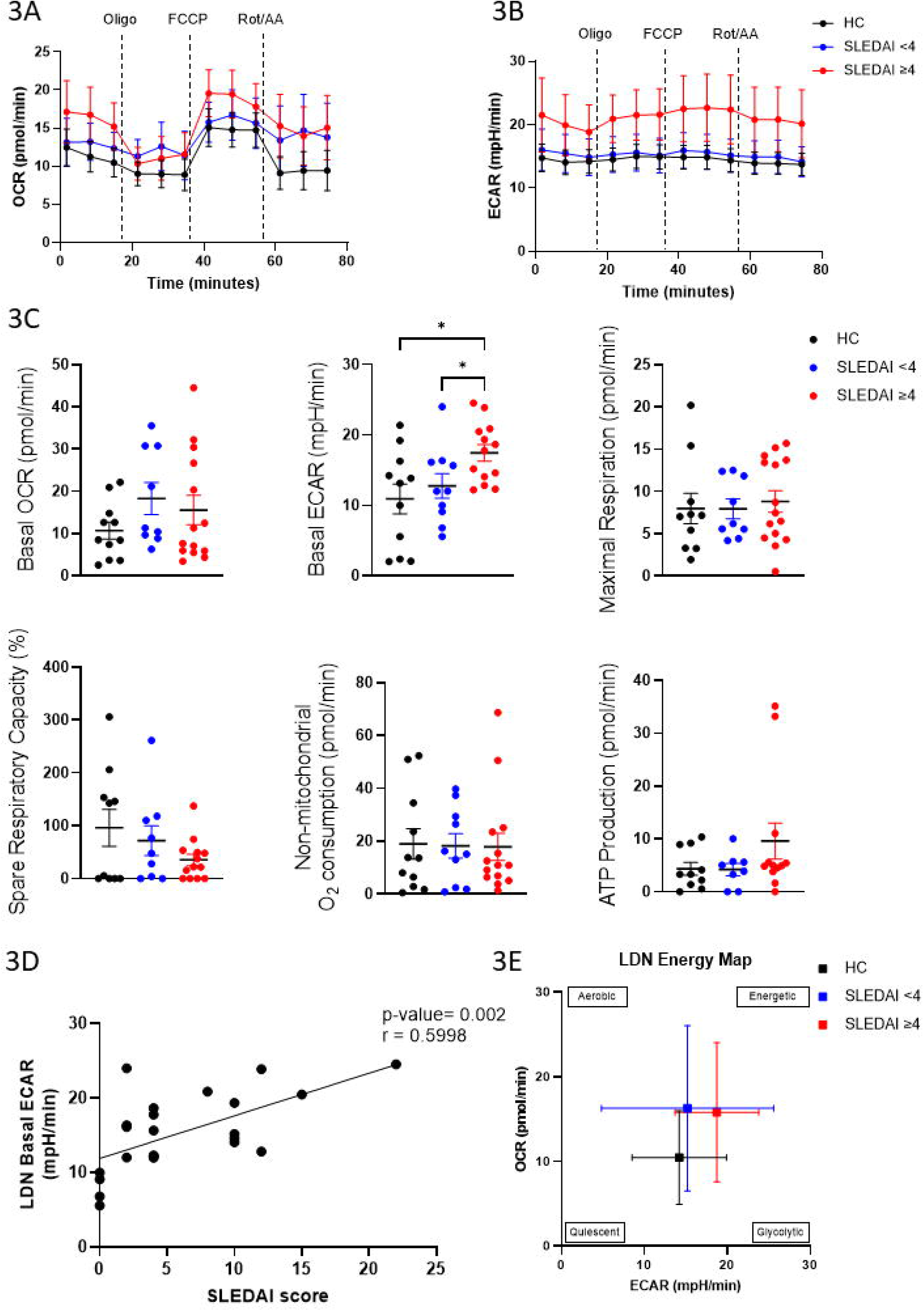
SLE LDNs exhibit increased glycolytic activity than HC LDNs, and this correlates with disease activity. (A) The Seahorse MitoStress test reveals that basal OCR kinetics are comparable across LDNs from SLE patients and HC. Both HC and SLE LDNs demonstrate sensitivity to FCCP indicating the use of mitochondrial respiration when bioenergetic demand is increased in LDNs as a whole. (B)LDNs from SLEDAI ≥4 patients appear to have higher basal ECAR kinetics compared to both SLEDAI <4 and HC LDNs, which appear to be comparable. (C) However, plotting individual values reveals increased basal glycolysis in SLE LDNs irrespective of disease activity levels. All other metabolic parameters have comparable values. (D) Plotting basal ECAR values against SLEDAI scores reveals a statistically significant, moderately high positive correlation. (E) The energy map shows the energetic profiles of SLE LDNs, relative the more metabolically quiescent HC LDNs at rest. *All graphs depict mean values with SEM, and all analyses were Kruskal-Wallis tests, *p<0*,*05. SLEDAI: SLE Disease Activity Index; OCR: Oxygen Consumption Rate; ECAR: Extracellular Acidification Rate; SEM: Standard Error of Mean*.

### SLE LDNs exhibit metabolic flexibility, phenotypic changes, and pro-inflammatory potential in response to metabolic modulation

Next, in order to assess the propensity of SLE LDNs to generate ATP in a metabolically stressful environment, we used oligomycin to block mitochondrial ATP production and thus commit the cell to intercellular ATP production from glycolysis. We found that blocking mitochondrial ATP production induced an increase in ECAR in LDNs from active SLE disease only, indicating metabolic flexibility in this sub-group (Fig 4A). This increase in glycolysis in LDNs from patients with SLEDAI ≥4 relative to LDNs from SLEDAI <4 patients and healthy individuals is also depicted in the energy map (Fig.4B). To investigate which steps of the glycolytic pathway might be associated with this phenomenon, we also assessed the expression of glycolysis-associated genes by qPCR (Fig.S2A). However, no statistically significant differences were noted in the expression of *HK2, PFKM, PKM2*, and *PFKFB3* between SLE LDNs with and without oligomycin treatment. This could potentially be explained by the short duration of oligomycin treatment, the difference in rate kinetics of metabolic and transcriptional machinery, or a potential need for mitochondrial metabolism in driving transcription, amongst other factors.

**Figure 4:**
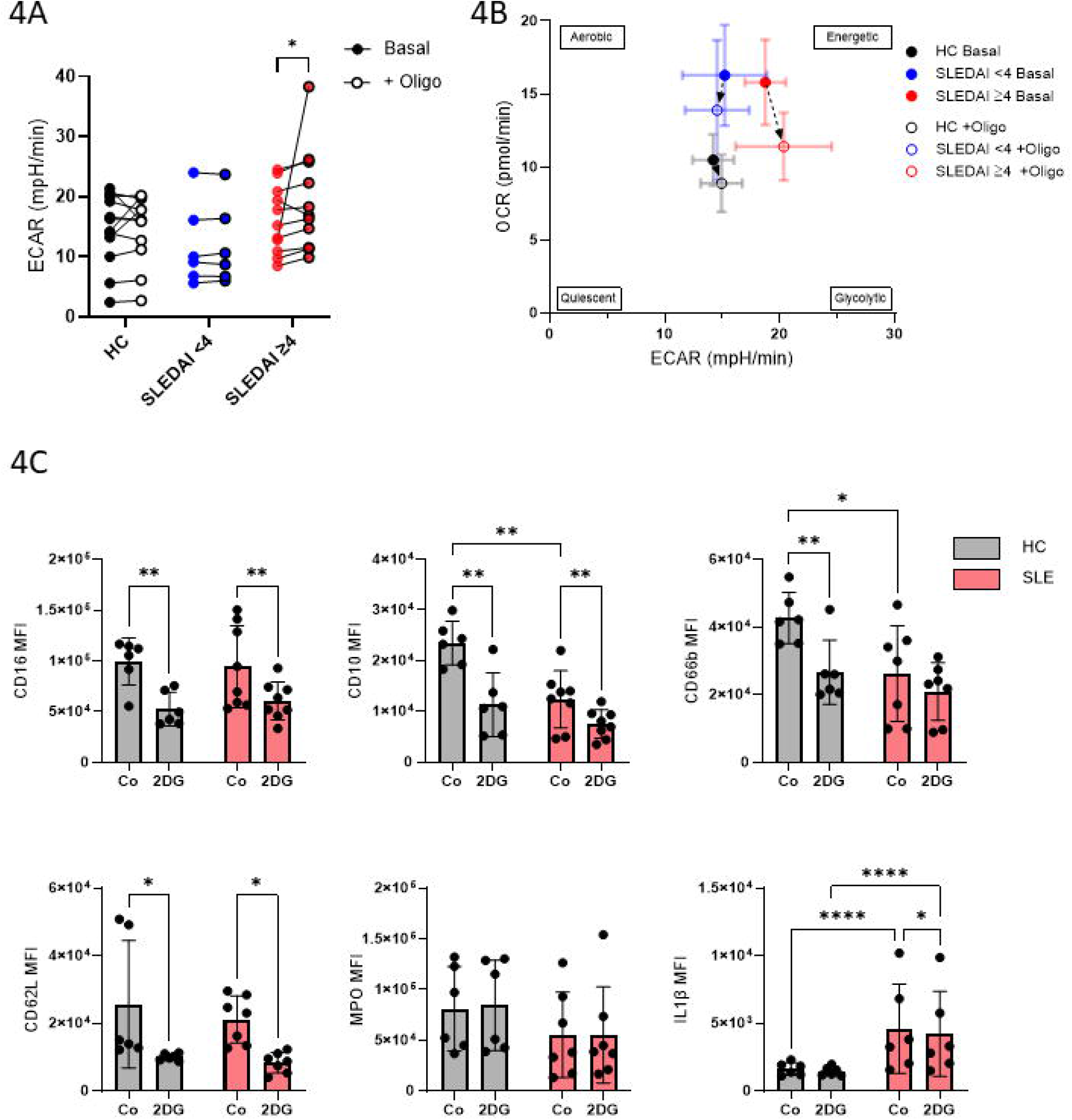
SLE LDNs exhibit metabolic flexibility, phenotypic changes, and pro-inflammatory potential in response to metabolic modulation. (A) The Seahorse MitoStress test showed that LDNs from SLEDAI ≥4 patients display an increase in glycolysis after mitochondrial ATPase is inhibited using oligomycin, and this is not seen in SLEDAI <4 or HC LDNs. (B) This mitochondrial flexibility of SLEDAI ≥4 LDNs is captured in the phenogram, through the shift to a relatively more glycolytic phenotype upon treatment with oligomycin. In comparison the relative shift in SLEDAI<4 and HC LDNs metabolism is negligible. (C) Bar plots show the change in MFI of common neutrophil markers of lineage maturation (CD10), activation (CD16, CD62L), and effector function (MPO, IE1β) at baseline (‘Co’) and after inhibition of glycolysis using 2DG. Graph (A) *depicts mean value, (B) depicts mean with SEM, (C) depicts and all analyses were Kruskal-Wallis tests, *p<0*,*05. SLEDAI: SLE Disease Activity Index; MFI: Median Fluorescence Intensity; 2DG: 2-deoxy-D-glucose; SEM: Standard Error of Mean*.

Since we observed that glycolytic activity is elevated in SLE LDNs relative to healthy LDNs, we propose that glycolysis may represent a viable therapeutic target. Accordingly, we sought to evaluate whether abrogation of excessive glycolysis in these cells using 2DG could alter their phenotype. For this, we evaluated the expression of identification (CD15, D66b), maturity (CD10), activation (CD16, CD62L), and effector/functional (MPO, IL-1β) markers in LDNs treated with 2DG (Fig.4C).

As paired analyses comparing SLE LDNs and NDNs by flow cytometry have already been explored in past studies and reveal no differences in MFIs, we focussed our assessments on SLE LDNs compared to HC LDNs. Matched analyses for the NDNs can be found in Figure S2C. We found that, in both healthy and SLE LDNs, inhibition of glycolysis caused a statistically significant reduction in the mean fluorescent intensity (MFI) of all markers except CD15, CD66b, and MPO (Fig.4C, S2B). We hypothesise that the reduction in phenotypic marker expression might be explained by the overall reduction of energetics brought on by glycolytic inhibition. CD10 MFI in SLE LDNs at baseline levels was lower compared to untreated HC LDNs, indicating an overall more immature population. This is in keeping with the compositional frequency of the LDN compartment in SLE (Fig.1E). Further, CD10 MFI reduced in both HC and SLE LDNs upon glycolytic inhibition. Interestingly, untreated LDNs from healthy individuals had intrinsically higher CD66b expression compared to SLE LDNs and were also responsive to 2DG treatment unlike SLE LDNs.

Finding that SLE LDNs engage in increased basal glycolysis, we expected a profile similar to activation and thus reduced CD62L expression at baseline. Counterintuitively, baseline CD62L expression in SLE LDNs was at par with HC LDNs. We also expected higher or comparable CD62L expression on treatment with 2DG but noted a reduction in both HC and SLE LDNs, which could potentially be explained by the cytotoxic effect of 2DG. IL-1β expression in SLE LDNs was found to be inherently higher compared to healthy LDNs independent of glycolytic inhibition, suggesting that SLE LDNs have higher levels of premade IL-1β. However, the marginal yet statistically significant reduction in IL-1β MFI after 2DG treatment indicates that, in SLE, LDN-derived IL-1β levels might be dependent on glycolysis, unlike in HC LDNs.

## Discussion

Since being described for the first time as a pro-inflammatory neutrophil subset 40 years ago, LDN research in SLE has largely focussed on defining their phenotype, effector function, and morphology ^1,19,26,26^. LDN immunometabolism in SLE has been largely overlooked, with less than a handful of studies focussed only on omics-based approaches ^3,22^, and no real-time metabolic studies reported to date. Additionally, metabolism-based studies in SLE and otherwise often assess neutrophils as a single unified population lacking distinction between subtypes. Our study sought to address these shortcomings and characterises neutrophil subsets in SLE with a specific focus on LDNs and their immunometabolism. We describe for the first time, that LDNs from SLE patients with increased disease severity exhibit elevated glycolysis compared to LDNs from healthy individuals. Notably, LDNs from these individuals also display metabolic flexibility and as such might play a distinct role in the pathogenesis of SLE.

Our work recapitulates findings of elevated and immature LDNs in SLE patients ^1,3,26,27^, and shows that the overall neutrophil landscape in patients with high disease severity is skewed toward a higher proportion of LDNs to NDNs, compared to healthy individuals. We have previously shown that chronic IFNα treatment *in vivo* induces elevated levels of immature granulocytes in circulation ^28^. Thus, we propose that the chronic inflammatory state in SLE driven by elevated type I IFN levels increases levels of immature LDNs in circulation. Other groups have defined differential roles of immune regulation for CD10 subsets in LDNs derived from GCSF-treated donors^2,3^ and in murine cancer models. Notably, CD10-LDNs produce increased NETs and ROS even in situations of glucose deprivation, indicating metabolic flexibility. This finding has implications for how this immature LDN subset might interact in microenvironments such as the vasculature or sites of skin lesions and contribute to SLE pathogenesis ^7,29^. Moreover, there has been a growing interest in utilising LDNs as biomarkers and measures of disease activity in SLE ^3,30,31^. The dichotomy in expression of markers like CD10, CD98, and CD177 indicates heterogeneity in the LDN compartment and has been proposed as a biomarker for disease progression in SLE ^2,3^. Our work extends the previously understood scope of heterogeneity within SLE LDNs and shows several sub-populations based on co-expression of neutrophil phenotypic markers.

In recent years, immunometabolism has become a pivotal concept in the field of immunology, inextricably linking cellular metabolism with immune cell function. In the present study, we use realtime metabolic techniques to show that in SLE patients with high disease activity, both LDNs and NDN use glycolysis at baseline more avidly compared to healthy neutrophils. Furthermore, basal glycolysis in LDNs from these patients was increased even in comparison to patients with low disease activity, indicating that the resting bioenergetic demand in SLE LDNs increases with disease severity. To our knowledge, no prior studies have directly examined *in situ* LDN metabolism in SLE. Therefore, to contextualize and support our findings, we drew on observations from a murine immature LDN study, which represents the closest available point of comparison to support our work. Hsu *et al* report that murine immature LDNs in glucose-abundant environments (such as in our study) engaged in glycolysis and not mitochondrial metabolism, and only an environment of glucose-deprivation caused a metabolic shift to mitochondrial respiration ^19^. NETosis, ROS production, and the release of pro-inflammatory cytokine production and bioenergetically demanding effector functions in neutrophils ^32,33^, and our finding of aberrantly elevated glycolysis, supported by that of Hsu et al, could explain the increased occurrence of NETs and release of pro-inflammatory cytokines associated with SLE LDNs reported in the literature ^1,4^.

In conducting our assessment of mitochondrial respiration in SLE LDNs, we observed that that inhibition of mitochondrial Complex V drove a compensatory increase in ECAR, unique only to patients with high disease activity and indicative of metabolic flexibility in LDNs from this cohort. Our observation that glycolytic activity is (i) elevated in LDNs from SLE patients with high disease activity at baseline, and (ii) induced as a feature of metabolic flexibility also suggests that, abrogating excessive glycolysis (for instance, using 2DG) in LDNs may represent a therapeutic strategy in this cohort. We propose that, if fine-tuned, metabolic inhibition could produce function-specific rewiring, and not necessarily a uniform shutdown of the cell’s metabolic machinery. Notably, immunophenotyping SLE LDNs and NDNs after glycolytic inhibition revealed the reduced expression of common markers for neutrophil maturity (CD10), activation and rolling (CD16, CD62L), and production of pro-inflammatory cytokines (IL-1β) implying that the expression of these fundamental markers is under glycolytic control. While our study focussed only on circulating neutrophils, our findings suggest that metabolic modulation of SLE LDNs have implications for the phenotypic alterations these cells might undergo at sites of tissue and vascular damage, as well as impact the interactions they have with other immune cells within these microenvironments.

Finally, although we assessed NDNs largely as an autologous control, our work provides insights into the metabolic profile of SLE NDNs. Like SLE LDNs, NDNs from the high disease activity cohort exhibit increased resting glycolysis. However, compared to LDNs, only SLE NDNs from patients with high disease activity exhibited mitochondrial involvement in response to increased metabolic demand. NDNs also underwent a similar reduction in phenotypic changes in response to glycolytic inhibition, however the pattern in reduction in SLE NDNs was different than that of SLE LDNs. These results combined are the first of their kind for SLE NDNs, highlighting a) a need for more studies examining immunometabolic differences between neutrophil subsets in SLE, and b) a difference in metabolic responsiveness compared to SLE LDNs that, if properly exploited, might aid in refining the use of LDNs as a therapeutic target.

### Limitations

Our results overall show correlations between disease severity and dysregulated bioenergetics in SLE LDNs and NDNs. However, a large number of patients in our cohort are on corticosteroid (~45%) and anti-malarial drugs (~80%). As disease severity and treatment are linked intrinsically, our results could be influenced by co-existent treatment with immunosuppressants or anti-malarial drugs. Neutrophil effector functions, detachment, and survival are shown to be affected by glucocorticoid treatment and anti-malarial drugs, but the exact effects have been a matter of debate^24,25^. Although very limited data exists on the effect of these drugs on cell metabolism, murine neutrophils treated with dexamethasone have been shown to increase glucose consumption but not glucose oxidation^25^, however no data in human neutrophils currently exists. Another study examining the effects of high-dose chloroquine on neutrophils was shown to alter oxidative metabolism^24^, however we do not report statistically significant changes in oxidative metabolism. While we did not investigate associations between our findings and these drugs, it is important to note that the impact of steroids could not be completely disregarded from this analysis.

In summary, our study provides the first real-time characterisation of immunometabolic activity of LDNs in SLE, identifying elevated glycolysis and enhanced metabolic flexibility as defining features of LDNs in patients with high disease activity. These findings expand the current understanding of LDN heterogeneity by linking metabolic state to functional phenotype and disease severity. Our data suggest that elevated bioenergetics, predominantly driven by glycolysis, may underpin key pathogenic functions of LDNs, including their pro-inflammatory and tissue-damaging potential. Importantly, the observed sensitivity of neutrophil phenotype to glycolytic inhibition highlights cellular metabolism as a potentially tractable axis for therapeutic intervention. Together, this work positions immunometabolism as a critical determinant of neutrophil behaviour in SLE and provides a foundation for future studies aimed at targeting metabolic pathways to modulate disease progression.

## Acknowledgements

We would like to thank Dr. Lorraine Thong, Dr. Kevin Brown, and Dr. Seán Donohue for their assistance in collecting samples from healthy donors for this manuscript.

---

**Supplementary Figure 1:**
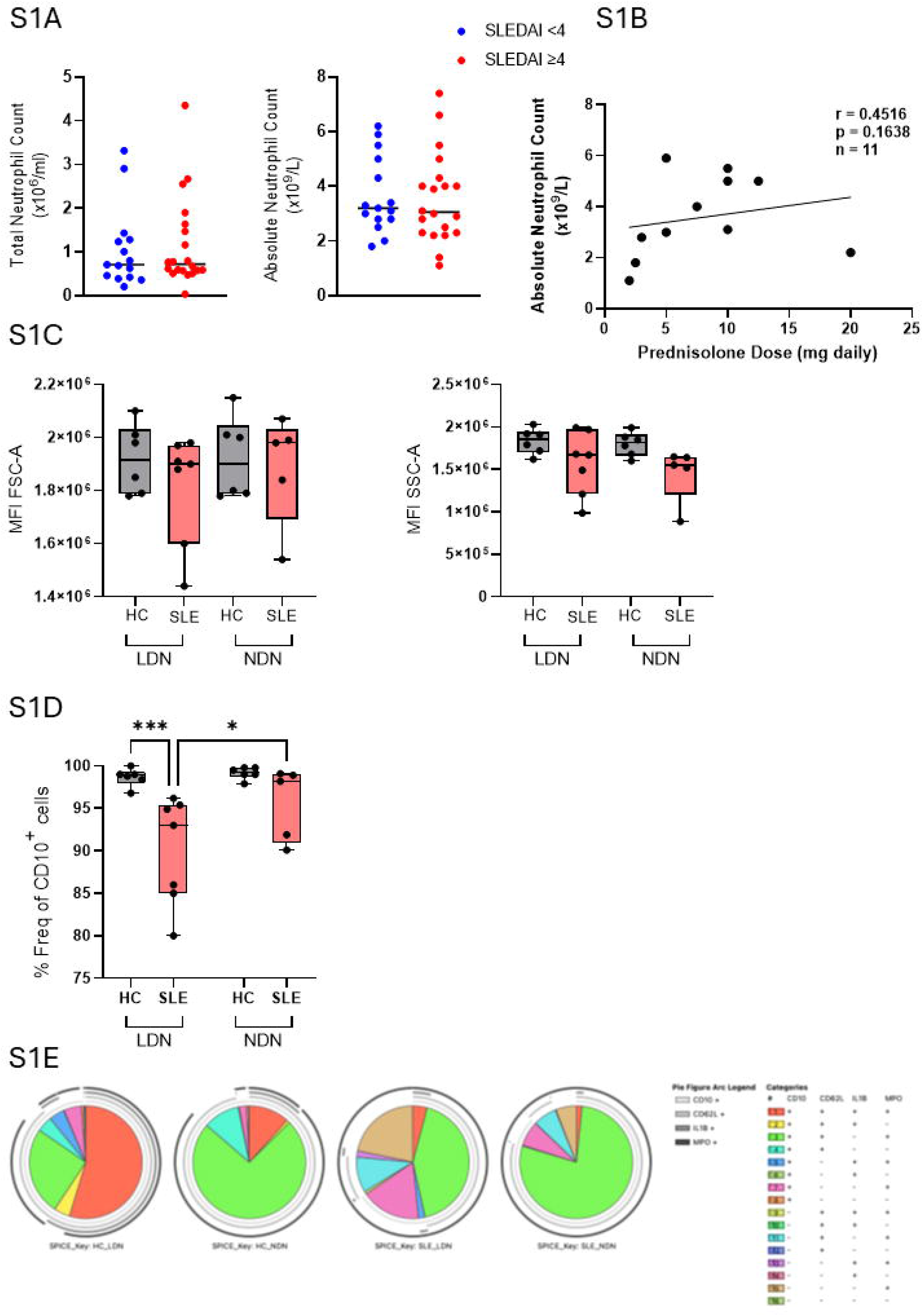
(A) Total neutrophil counts obtained after magnetic isolation, and ANC obtained from clinical reports are comparable between SLEDAI ≥4 and SLEDAI <4 patients. (B) Clinically obtained ANC plotted against Prednisolone dose do not show statistically significant correlation between prescribed steroid dose and circulating neutrophil counts. (B) FSC-A and SSC-A MFIs reveal no statistically significant differences in cell size or granularity, respectively, across HC and SLE neutrophil subsets. (C) SLE patients have a reduced frequency of mature (CD10+) LDNs compared to HC LDNs and autologous NDNs. (D) Pie charts generated following SPICE analysis subpopulation frequencies indicate heterogeneity in SLE LDNs, defined by differences in co-expression of CD10, CD62L, IL1β, and MPO. *All analyses were Kruskal-Wallis tests, *p<0*,*05. ANC: Absolute Neutrophil Count; SLEDAI: SLE Disease Activity Index; MFI: Median Fluorescent Intensity; FSC-A: Forward Scatter Area; SSC-A: Side Scatter Area; SPICE: Simplified Presentation of Incredibly Complex Evaluations*.

**Supplementary Figure 2:**
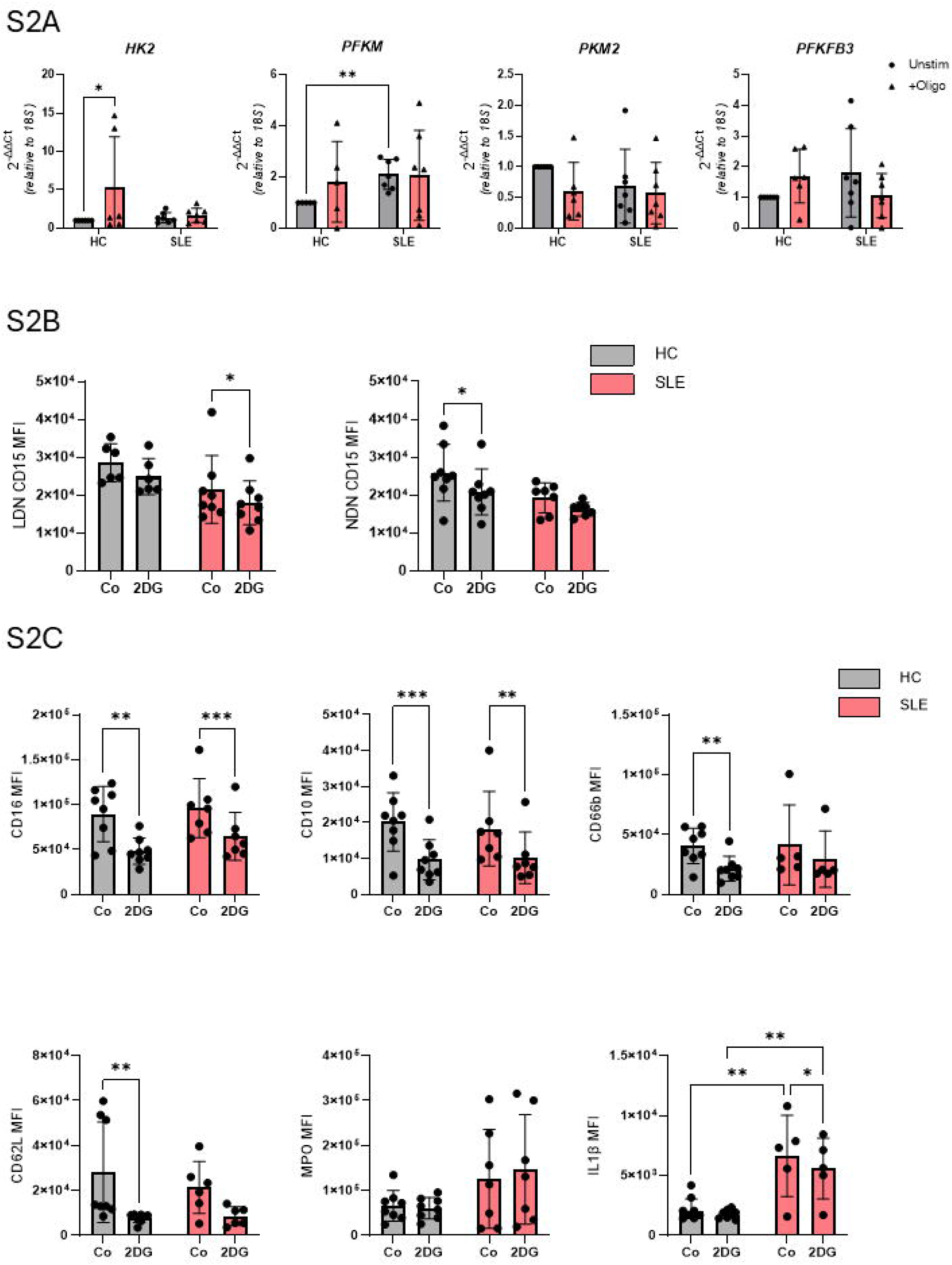
(A) qPCR assessment of glycolytic genes in LDNs revealed no statistically significant difference between SLE LDNs upon treatment with oligomycin to inhibit ATPase. (B) Changes in LDN and NDN CD15 MFI after inhibition of glycolysis using 2DG. (C) Bar plots show the change in NDN MFI of common neutrophil markers of lineage maturation (CD10), activation (CD16, CD62L), and effector function (MPO, IL1β) at baseline (‘Co’) and after inhibition of glycolysis using 2DG. *All analyses were Kruskal-Wallis tests, *p<0*,*05. MFI: Median Fluorescence Intensity; 2DG: 2-deoxy-D-glucose*.

## References

1 Denny, M. F. et al. A distinct subset of proinflammatory neutrophils isolated from patients with systemic lupus erythematosus induces vascular damage and synthesizes type I IFNs. Journal of immunology (Baltimore, Md.:1950) 184, 3284–3297 (2010). 10.4049/jimmunol.0902199

2 Marini, O. et al. Mature CD10(+) and immature CD10(-) neutrophils present in G-CSF-treated donors display opposite effects on T cells. Blood 129, 1343–1356 (2017). 10.1182/blood-2016-04-713206

3 Martin, K. R. et al. CD98 defines a metabolically flexible, proinflammatory subset of low-density neutrophils in systemic lupus erythematosus. Clinical and Translational Medicine 13, e1150 (2023). 10.1002/ctm2.1150

4 Villanueva, E. et al. Netting neutrophils induce endothelial damage, infiltrate tissues, and expose immunostimulatory molecules in systemic lupus erythematosus. Journal of immunology (Baltimore, Md.:1950) 187, 538–552 (2011). 10.4049/jimmunol.1100450

5 Bashant, K. R. et al. Proteomic, biomechanical and functional analyses define neutrophil heterogeneity in systemic lupus erythematosus. Ann Rheum Dis 80, 209–218 (2021). 10.1136/annrheumdis-2020-218338

6 Aderka, D. et al. Correlation between serum levels of soluble tumor necrosis factor receptor and disease activity in systemic lupus erythematosus. Arthritis Rheum 36, 1111–1120 (1993). 10.1002/art.1780360812

7 Yennemadi, A. S., Jordan, N., Diong, S., Keane, J. & Leisching, G. The Link Between Dysregulated Immunometabolism and Vascular Damage: Implications for the Development of Atherosclerosis in Systemic Lupus Erythematosus and Other Rheumatic Diseases. J Rheumatol 51, 234–241 (2024). 10.3899/jrheum.2023-0833

8 Mistry, P. et al. Transcriptomic, epigenetic, and functional analyses implicate neutrophil diversity in the pathogenesis of systemic lupus erythematosus. Proceedings of the National Academy of Sciences 116, 25222–25228 (2019).

9 Yennemadi, A. S., Keane, J. & Leisching, G. The Isolation and Characterization of Low-and Normal-Density Neutrophils from Whole Blood. J Vis Exp (2025). 10.3791/67805

10 Yennemadi, A. S., Murphy, F. K., Keane, J. & Leisching, G. Sex-specific metabolic programming in human neutrophil subsets. J Leukoc Biol 118 (2025). 10.1093/jleuko/qiaf179

11 Hardisty, G. R. et al. High Purity Isolation of Low Density Neutrophils Casts Doubt on Their Exceptionality in Health and Disease. Frontiers in Immunology 12 (2021). 10.3389/fimmu.2021.625922

12 Jeon, J. H., Hong, C. W., Kim, E. Y. & Lee, J. M. Current Understanding on the Metabolism of Neutrophils. Immune Netw 20, e46 (2020). 10.4110/in.2020.20.e46

13 Aringer, M. et al. 2019 European League Against Rheumatism/American College of Rheumatology classification criteria for systemic lupus erythematosus. Arthritis & rheumatology 71, 1400–1412 (2019).

14 Roederer, M., Nozzi, J. L. & Nason, M. C. SPICE: exploration and analysis of post-cytometric complex multivariate datasets. Cytometry A 79, 167–174 (2011). 10.1002/cyto.a.21015

15 Schmittgen, T. D. & Livak, K. J. Analyzing real-time PCR data by the comparative C(T) method. Nat Protoc 3, 1101–1108 (2008). 10.1038/nprot.2008.73

16 Evrard, M. et al. Developmental Analysis of Bone Marrow Neutrophils Reveals Populations Specialized in Expansion, Trafficking, and Effector Functions. Immunity 48, 364–379 e368 (2018). 10.1016/j.immuni.2018.02.002

17 Morosetti, R. et al. A novel, myeloid transcription factor, C/EBP epsilon, is upregulated during granulocytic, but not monocytic, differentiation. Blood 90, 2591–2600 (1997).

18 Yamanaka, R. et al. CCAAT/enhancer binding protein epsilon is preferentially up-regulated during granulocytic differentiation and its functional versatility is determined by alternative use of promoters and differential splicing. Proc Natl Acad Sci U S A 94, 6462–6467 (1997). 10.1073/pnas.94.12.6462

19 Hsu, B. E. et al. Immature Low-Density Neutrophils Exhibit Metabolic Flexibility that Facilitates Breast Cancer Liver Metastasis. Cell Rep 27, 3902–3915 e3906 (2019). 10.1016/j.celrep.2019.05.091

20 Hassani, M. et al. On the origin of low-density neutrophils. J Leukoc Biol 107, 809–818 (2020). 10.1002/JLB.5HR0120-459R

21 Parackova, Z. et al. Expanded population of low-density neutrophils in juvenile idiopathic arthritis. Front Immunol 14, 1229520 (2023). 10.3389/fimmu.2023.1229520

22 Tay, S. H., Celhar, T. & Fairhurst, A.-M. Low-Density Neutrophils in Systemic Lupus Erythematosus. Arthritis & Rheumatology 72, 1587–1595 (2020). 10.1002/art.41395

23 Chacko, B. K. et al. Methods for defining distinct bioenergetic profiles in platelets, lymphocytes, monocytes, and neutrophils, and the oxidative burst from human blood. Lab Invest 93, 690–700 (2013). 10.1038/labinvest.2013.53

24 Labro, M. T. & Babin-Chevaye, C. Effects of amodiaquine, chloroquine, and mefloquine on human polymorphonuclear neutrophil function in vitro. Antimicrob Agents Chemother 32, 1124–1130 (1988). 10.1128/AAC.32.8.1124

25 Garcia, C., de Oliveira, M. C., Verlengia, R., Curi, R. & Pithon-Curi, T. C. Effect of dexamethasone on neutrophil metabolism. Cell Biochem Funct 21, 105–111 (2003). 10.1002/cbf.1002

26 Rahman, S. et al. Low-density granulocytes activate T cells and demonstrate a non-suppressive role in systemic lupus erythematosus. Ann Rheum Dis 78, 957–966 (2019). 10.1136/annrheumdis-2018-214620

27 Bennett, L. et al. Interferon and granulopoiesis signatures in systemic lupus erythematosus blood. J Exp Med 197, 711–723 (2003). 10.1084/jem.20021553

28 Yennemadi, A. S. et al. Chronic IFNα treatment induces leukopoiesis, increased plasma succinate and immune cell metabolic rewiring. Cellular Immunology 390, 104741 (2023). 10.1016/j.cellimm.2023.104741

29 Safi, R., Al-Hage, J., Abbas, O., Kibbi, A. G. & Nassar, D. Investigating the presence of neutrophil extracellular traps in cutaneous lesions of different subtypes of lupus erythematosus. Exp Dermatol 28, 1348–1352 (2019). 10.1111/exd.14040

30 Xu, H. et al. Aberrant expansion of CD177(+) neutrophils promotes endothelial dysfunction in systemic lupus erythematosus via neutrophil extracellular traps. J Autoimmun 152, 103399 (2025). 10.1016/j.jaut.2025.103399

31 Henning, S. et al. Low-density granulocytes and neutrophil extracellular trap formation are increased in incomplete systemic lupus erythematosus. Rheumatology (Oxford) 64, 1234–1242 (2025). 10.1093/rheumatology/keae300

32 Freitag, A. et al. Glucose metabolism controls oxidative burst and lipid mediator production in neutrophils upon microbial challenge. Microlife 6, uqaf040 (2025). 10.1093/femsml/uqaf040

33 Kumar, S. & Dikshit, M. Metabolic Insight of Neutrophils in Health and Disease. Front Immunol 10, 2099 (2019). 10.3389/fimmu.2019.02099

